# Targeting RTN4/NoGo-Receptor reduces levels of ALS protein ataxin-2

**DOI:** 10.1101/2021.12.20.473562

**Authors:** Caitlin M. Rodriguez, Sophia C. Bechek, Graham L. Jones, Lisa Nakayama, Tetsuya Akiyama, Garam Kim, David E. Solow-Cordero, Stephen M. Strittmatter, Aaron D. Gitler

## Abstract

Gene-based therapeutic strategies to lower ataxin-2 levels are emerging for neurodegenerative diseases amyotrophic lateral sclerosis (ALS) and spinocerebellar ataxia type 2 (SCA2). To identify additional ways of reducing ataxin-2 levels, we performed a genome-wide screen in human cells for regulators of ataxin-2 and identified *RTN4R*, the gene encoding the RTN4/NoGo-Receptor, as a top hit. *RTN4R* knockdown, or treatment with a peptide inhibitor, was sufficient to lower ataxin-2 protein levels in mouse and human neurons *in vitro* and *Rtn4r* knockout mice have reduced ataxin-2 levels *in vivo*. Remarkably, we observed that ataxin-2 shares a role with the RTN4/NoGo-Receptor in limiting axonal regeneration. Reduction of either protein increases axonal regrowth following axotomy. These data define the RTN4/NoGo-Receptor as a novel therapeutic target for ALS and SCA2 and implicate the targeting of ataxin-2 as a potential treatment following nerve injury.

## Introduction

ALS is a rapidly progressive neurodegenerative disease characterized by loss of motor neurons from the brain and spinal cord. Loss of motor neurons leads to muscle weakness, paralysis, and eventual death 3-5 years following diagnosis (Taylor et al., 2016). SCA2 is associated with loss of Purkinje neurons from the cerebellum, which affects balance and coordination, and causes slow saccadic eye movements and cognitive impairment (Paulson et al., 2017). SCA2 is an autosomal dominant genetic disorder caused by long CAG repeat expansions (>34) in the *ATXN2* gene, which encodes the ataxin-2 protein (Scoles and Pulst, 2018). Alternatively, intermediate length repeat expansions (27-33) in *ATXN2* are a genetic risk factor for ALS (Elden et al., 2010). Therapeutic strategies to target ataxin-2 have shown efficacy in preclinical studies (Becker et al., 2017, Scoles et al., 2017), motivating the initiation of a clinical trial testing *ATXN2-*targeting antisense oligonucleotides (ASOs) in human ALS patients (ClinicalTrials.gov identifier: NCT04494256).

## Results

### Whole-genome siRNA screen identifies genes and cellular pathways that regulate ataxin-2 levels

To discover additional targets and pathways to lower ataxin-2 levels, we created a quantitative and highly sensitive reporter of endogenous ataxin-2 protein levels by inserting a HiBiT tag on the C-terminus of ataxin-2 in HEK293T cells using CRISPR-Cas9 genome editing (**Figure 1A**). HiBiT is an 11-amino acid peptide subunit of the NanoBiT luciferase enzyme that is inert on its own but binds with high affinity to LgBiT, reconstituting NanoBiT (Dixon et al., 2016). This allows for easy and specific detection of ataxin-2 using an antibody-free blotting method (**Figure 1B-D**). An in-well reaction of ataxin-2-HiBiT cell lysate with the exogenous addition of LgBiT and the furimazine substrate generates quantifiable bioluminescence as a direct measure of ataxin-2 abundance (**Figure 1E**). In parallel, we stably incorporated firefly luciferase (FFluc) into the ataxin-2-HiBiT line to normalize for protein abundance.

**Figure 1:**
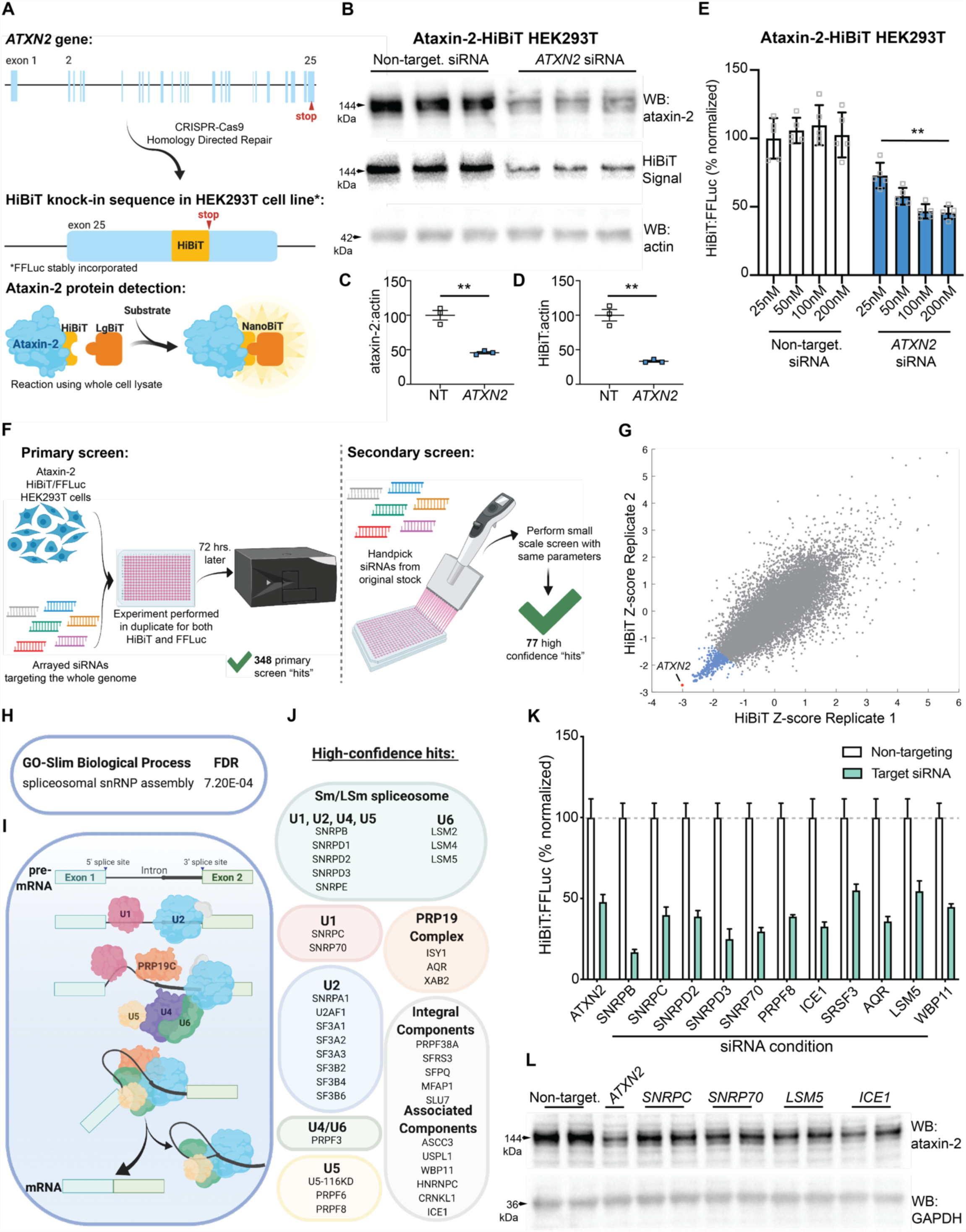
Generation of the ataxin-2-HiBiT cell line and overview of the whole-genome siRNA screen. **(A)** We engineered an endogenous C-terminal HiBiT fusion on ataxin-2 using CRISPR-Cas9 genome editing in HEK293T cells. LgBiT compliments the HiBiT protein tag to form NanoBiT. As a control, we stably incorporated firefly luciferase (FFLuc) into this line. **(B)** Antibody-based immunoblotting and HiBiT substrate-based detection on ataxin-2-HiBiT cell lysates transfected with siRNA. Quantified in **(C)** and **(D). (E)** HiBiT signal measured via luciferase assay on cells transfected with increasing doses of siRNA. **(F)** Schematic of the whole-genome siRNA screen for regulators of ataxin-2 levels. **(G)** Plot showing results of HiBiT replicates after filtering for changes in FFLuc. Datapoints in blue are primary screen hits that decrease ataxin-2. *ATXN2* siRNA (red) was the strongest hit. **(H)** GO-Slim Biological Process analysis of the hits from the primary screen. **(I)** Summary of the major steps in pre-mRNA splicing, illustrating the involvement of the PRP19-associated complex (PRP19C) and the U1, U2, U4, U5, and U6 snRNP RNA-protein complexes. **(J)** A list of the siRNAs that reduce ataxin-2 validated in the secondary screen. **(K)** Ataxin-2-HiBiT cells treated with siRNAs against several splicing components followed by a luciferase assay to measure HiBiT activity. Each splicing factor knockdown significantly reduces HiBiT signal relative to non-targeting. Two-way ANOVA with multiple comparisons: *WBP11, SRSF3*, and *LSM5*, p ≤ 0.001. *ATXN2, SNRPB, SNRPC, SNRPD2, SNRPD3, SNRP70, PRPF8, ICE1*, and *AQR*, p ≤ 0.0001. **(L)** Immunoblot of unedited HEK293T cell lysates after siRNA treatment. C and D, Student’s t-test. E, One-way ANOVA with multiple comparisons. **p ≤ 0.01. Error bars represent ± SEM.

We used the ataxin-2-HiBiT cell line to perform a genome-wide screen using arrayed siRNA pools targeting 21,121 mRNA transcripts (**Figure 1F, 1G, and S1**). We performed the screen in duplicate for both ataxin-2-HiBiT levels and FFluc levels, a control for ruling out siRNAs with nonspecific effects on gene expression. We classified siRNAs that had an average HiBiT Z-score less than −1.65 and an average FFluc Z-score greater than −1 as a hit that decreased ataxin-2— resulting in 348 primary screen hits (**Supplementary Data Table 1**). We selected primary screen hits for validation based on function in shared pathways (e.g., splicing) or known roles in the nervous system. We performed a secondary validation screen using 102 siRNA pools, of which 77 validated and we classified these as “high confidence hits” (**Figure S1C, Table S1**).

Gene ontology analysis of the primary screen hits revealed an enrichment in constituents of the pre-mRNA splicing pathway (**Figure 1H and 1I**), and 36 of the 77 high confidence hits have roles in the major spliceosome complexes (**Figure 1J**). These proteins are enriched for the like-Sm or LSm domain (**Figure S2**), a protein domain critical for complex formation and splicing activity (He and Parker, 2000). Ataxin-2 has an LSm domain that is one of the few predicted structured regions of this largely disordered protein. We independently validated several high confidence hits using siRNAs in unedited HEK293T cells and immunoblotting for ataxin-2 (**Figure 1K, 1L, and S3A**). We also performed RT-qPCR and found that some of these hits affected steady-state *ATXN2* mRNA levels (**Figure S3B**). The presence of a shared LSm domain combined with the regulation revealed by this screen suggests a potential functional connection between ataxin-2 and the spliceosome.

### Knockdown or inhibition of the RTN4/NoGo-Receptor lowers ataxin-2 levels

Because splicing is an essential cellular process and splicing defects contribute to ALS (Lagier-Tourenne et al., 2010), we sought to identify the optimal target with therapeutic potential by filtering high confidence hits for essentiality (DepMap database, average gene effect>-1) (Meyers et al., 2017, McFarland et al., 2018) and central nervous system expression (GTEx database, cortical pTPM>15) (2013) (**Figure 2A**). We reasoned that selecting targets with high nervous system specificity may eliminate potential negative off-target effects in non-diseased tissues. Following these filtering steps, the top hit was *RTN4R*, the gene that encodes RTN4/NoGo-Receptor. RTN4/NoGo-Receptor has been implicated in axon regeneration, sprouting, and plasticity (Fournier et al., 2001, McGee et al., 2005, Wang et al., 2020, Wang et al., 2011, Akbik et al., 2013, Bhagat et al., 2016, Kim et al., 2004, Fink et al., 2015), and targeting its ligand NoGo-A modulates mutant SOD1 mouse models of ALS (Jokic et al., 2006, Bros-Facer et al., 2014, Fournier et al., 2001, Yang et al., 2009). RTN4/NoGo-Receptor has several glia- and neuron-derived ligands including the three gene products of the *RTN4* gene—NoGo-A, B, and C—as well as oligodendrocyte myelin glycoprotein (OMgp) and myelin-associated glycoprotein (MAG) (Schwab, 2010, Domeniconi et al., 2002, Wang et al., 2002). Other high-affinity ligands include BAI adhesion-GPCRs (Wang et al., Chong et al., 2018), LGI1 (Thomas et al., 2010), BLyS (Zhang et al., 2009), and LOTUS (Sato et al., 2011). *RTN4R* knockdown reduced ataxin-2 levels by HiBiT assay and immunoblot (**Figure 2B-D**). We confirmed this in SH-SY5Y neuroblastoma cells (**Figure S4A and S4B**). Conversely, knockdown of ataxin-2 had no effect on RTN4/NoGo-Receptor levels (**Figure S4C and S4D**). This effect seems specific to ataxin-2 and not polyQ proteins in general because knocking down *RTN4R* did not affect expression levels of polyQ disease proteins huntingtin and ataxin-3 nor the ataxin-2 paralog ATXN2L (**Figure S5**). As another functional readout of decreased ataxin-2 function (Becker et al., 2017), *RTN4R* knockdown reduced recruitment of TDP-43 to stress granules (**Figure S6**). These data provide evidence that RTN4/NoGo-Receptor is required to maintain ataxin-2 levels.

**Figure 2:**
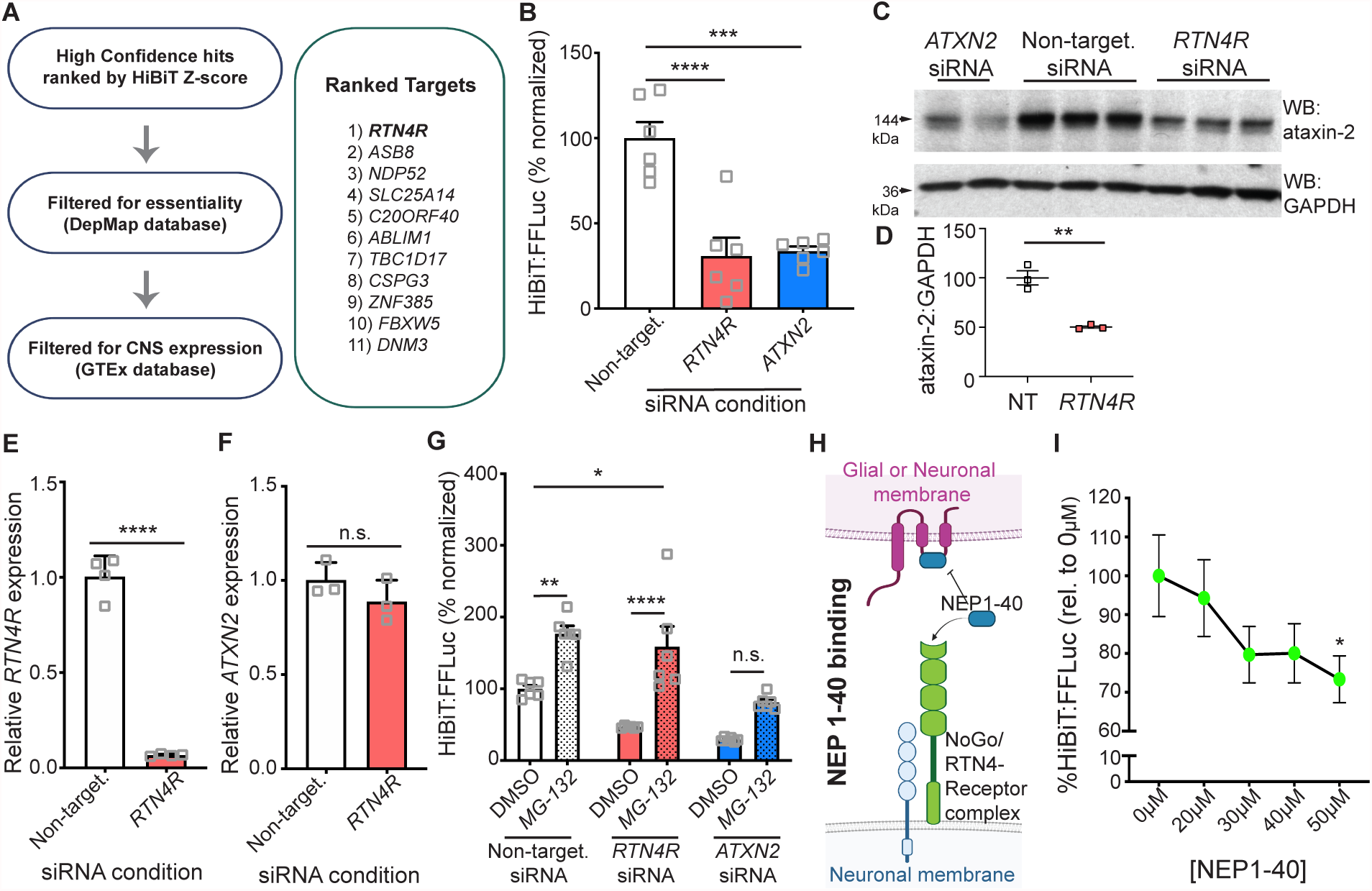
Targeting *RTN4R* lowers levels of ataxin-2. **(A)** High confidence hits ranked by average HiBiT Z-score, then filtered for essentiality (gene effect, DepMap) and CNS expression (GTEx). **(B)** HiBiT signal measured by luciferase assay in ataxin-2-HiBiT cells transfected with designated siRNA. **(C)** Immunoblot for ataxin-2 and GAPDH on lysates derived from unedited HEK293T cells treated with designated siRNA. Ataxin-2 levels are quantified in **(D)**. We performed RT-qPCR on RNA from cells treated as in (C). We probed for *RTN4R* transcript **(E)** or *ATXN2* transcript **(F)** along with *ACTB* for normalization. **(G)** We treated ataxin-2-HiBiT cells with siRNA then treated cells for 8hr with proteasome inhibitor MG-132 or DMSO and performed a luciferase assay to measure HiBiT activity. **(H)** The NEP1-40 peptide, a shared extracellular region of the NoGo proteins, binds to RTN4/NoGo-Receptor and prevents further signaling through the receptor. **(I)** We treated ataxin-2-HiBiT cells for 48hr with increasing doses of NEP1-40 and performed a luciferase assay to measure HiBiT activity. B and I, One-way ANOVA with multiple comparisons. G, Two-way ANOVA with multiple comparisons. D, E, and F, Student’s t-test. *p ≤ 0.05, **p ≤ 0.01, ***p ≤ 0.001, ****p ≤ 0.0001. Error bars represent ± SEM.

Regulation of ataxin-2 by RTN4/NoGo-Receptor occurs at the protein level because *RTN4R* knockdown did not affect *ATXN2* mRNA levels (**Figure 2E and 2F**). The HiBiT system allows for monitoring protein degradation (Riching et al., 2018). We used the ataxin-2-HiBiT line to test if *RTN4R* knockdown leads to ataxin-2 protein degradation by the proteasome or autophagy. Following siRNA knockdown of *RTN4R*, treatment of cells with a proteasome inhibitor (**Figure 2G)**, but not an autophagy inhibitor (**Figure S7A)**, resulted in an increase in ataxin-2 to levels comparable to the non-targeting control, indicating that ataxin-2 is degraded by the proteasome. *RTN4R* knockdown did not increase 20S proteasome activity (**Figure S7B**), suggesting a specific effect on ataxin-2 proteasomal degradation caused by *RTN4R* knockdown.

NEP1-40 is a peptide that acts as a competitive RTN4/NoGo-Receptor antagonist. It is a fragment of the luminal region of NoGo-A, B and C that binds to RTN4/NoGo-Receptor to prevent ligand signaling (**Figure 2H**) (GrandPré et al., 2002). NEP1-40 treatment decreased ataxin-2 levels in the ataxin-2-HiBiT cells (**Figure 2I**). Thus, targeting *RTN4R* by either genetic knockdown or with a peptide inhibitor leads to decreased levels of ataxin-2.

### Knockdown or inhibition of the RTN4/NoGo-Receptor lowers ataxin-2 levels in mouse and human neurons

To extend our findings to neurons, we tested this interaction in mouse cortical neurons and human iPSC-derived neurons (iNeurons) (**Figure 3A and 3B**) (Bieri et al., 2019). Treatment of mouse cortical neurons and human iNeurons with lentiviruses expressing shRNA targeting mouse or human *RTN4R* respectively resulted in about a 50% reduction of ataxin-2 (**Figure 3C-F**), without affecting levels of *ATXN2* mRNA (**Figure S8**). Application of the NEP1-40 inhibitor peptide caused a dose-dependent reduction of ataxin-2 in both mouse cortical neurons and human iNeurons (**Figure 3G-L**). These results suggest that targeting RTN4/NoGo-Receptor, either genetically or with a peptide inhibitor, is sufficient to markedly decrease levels of ataxin-2 in neurons. Finally, we tested the effects of RTN4/NoGo-Receptor on ataxin-2 levels *in vivo*. We analyzed heterozygous and homozygous *Rtn4r* knockout mice. These mice are fertile and viable with no apparent deficits to the nervous system (Kim et al., 2004). We observed a dose-dependent reduction in ataxin-2 in lysates from mouse cortex from the *Rtn4r* heterozygous and homozygous knockout mice compared to wild-type mice (**Figure 3M and 3N**), showing that reduction of the RTN4/NoGo-Receptor is sufficient to lower levels of ataxin-2 in the nervous system.

**Figure 3:**
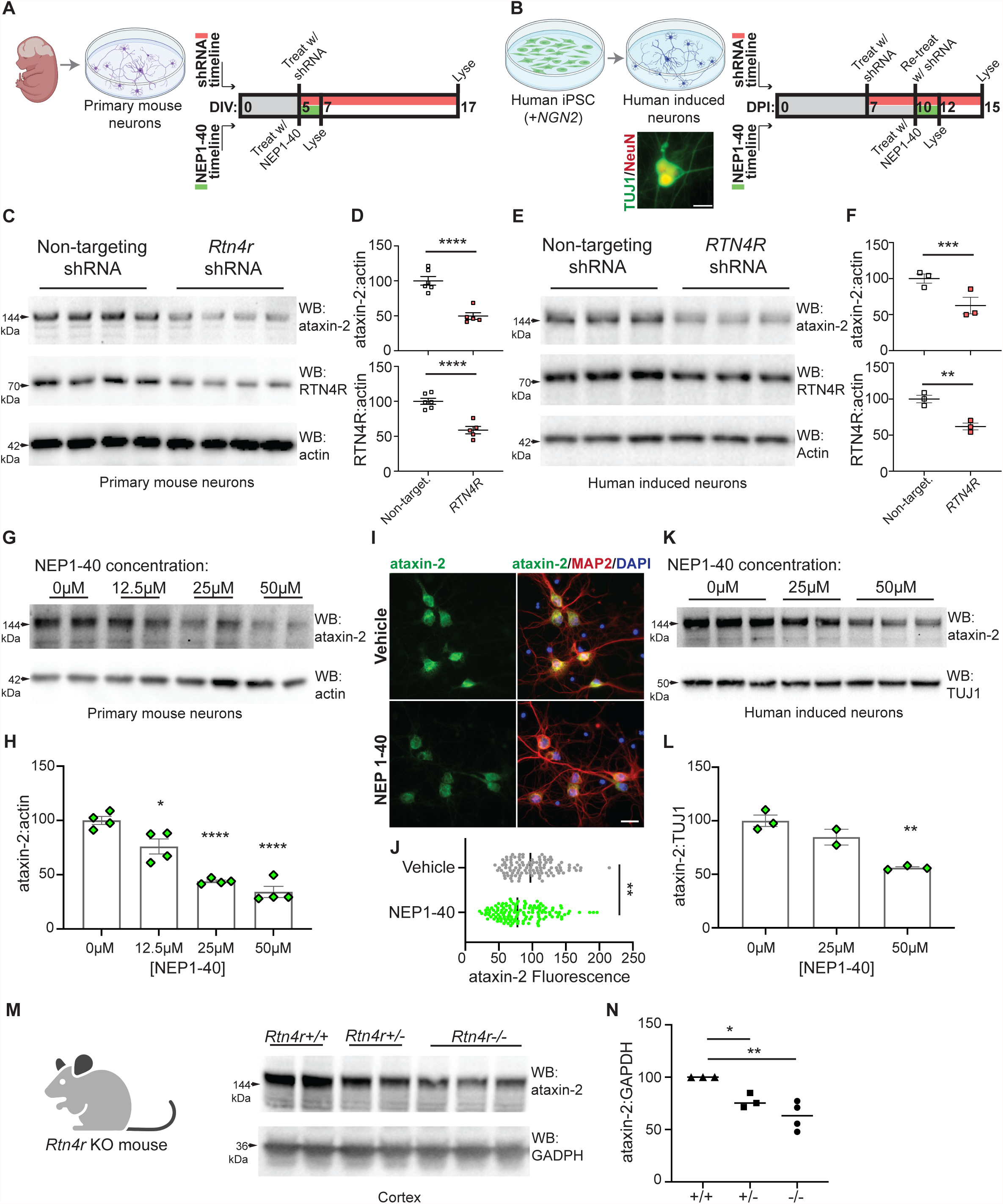
*RTN4R* knockdown or inhibition of RTN4/NoGo-Receptor in mouse and human neurons and in mouse brain reduces ataxin-2 levels. **(A)** Primary neurons isolated from embryonic mouse cortex. Treatment timeline for shRNA lentivirus or RTN4/NoGo-Receptor peptide inhibitor NEP1-40 in primary mouse neurons. DIV=days *in vitro*. **(B)** Induced neuron differentiation in a human iPSC line with *NGN2* stably integrated, as verified by Tuj1 (green) and NeuN (red) immunostaining, scale bar 20μm. Treatment timeline for shRNA lentivirus or NEP1-40 in human induced neurons (iNeurons). DPI=days post induction. **(C)** Immunoblot on lysates from mouse neurons treated with shRNA. **(D)** Quantification of ataxin-2 and RTN4/NoGo-Receptor levels. **(E)** Immunoblot on lysates from iNeurons treated with shRNA. **(F)** Quantification of ataxin-2 and RTN4/NoGo-Receptor. **(G)** Immunoblot on lysates from primary mouse neurons treated with increasing doses of NEP1-40. Quantification is shown in **(H). (I)** Immunocytochemistry and fluorescence microscopy on mouse neurons treated with 50 μM NEP1-40. MAP2 labels neurons, DAPI labels nuclei. Scale bar=20μm. **(J)** Quantification of neuronal ataxin-2 fluorescence. **(K)** Immunoblot on lysates from iNeurons treated for 48 hours with increasing doses of NEP1-40. Quantification is shown in **(L). (M)** Immunoblot on whole cortex lysates from *RTNR* +/+, +/-, and -/- mice. Quantification in **(N)**. D, F, and J, Student’s t-test. H, L, and N, One-way ANOVA with multiple comparisons. *p ≤ 0.05, **p ≤ 0.01, ***p ≤ 0.001, ****p ≤ 0.0001. Error bars represent ± SEM.

### Knockdown of ataxin-2 improves axon regrowth after injury

RTN4/NoGo-Receptor signaling destabilizes the actin cytoskeleton leading to growth cone collapse, limiting neurite outgrowth (Chivatakarn et al., 2007, Schwab, 2010, Montani et al., 2009). Reduction or inhibition of RTN4/NoGo-Receptor or its ligands has been shown to limit axonal regeneration following injury (Wang et al., 2020, Wang et al., 2011, Kim et al., 2004, Fink et al., 2015, Fournier et al., 2001, Schwab and Strittmatter, 2014). Since we have demonstrated that knocking down or inhibiting RTN4/NoGo-Receptor lowers ataxin-2, we tested if reducing ataxin-2 levels is sufficient to promote axon regeneration, perhaps functioning downstream of RTN4/NoGo-Receptor. We plated primary mouse neurons in the soma compartment of microfluidics chambers (**Figure 4A**), treated with lentivirus expressing shRNA targeting either *Atxn2* or *Rtn4r* at DIV5 (**Figure 4B**), and allowed neurons to mature and project axons through the microchannels and into the inner chamber of the axonal compartment. At DIV17, we performed vacuum-assisted axotomy to fully sever axons projecting into the inner chamber (**Figure 4C**) and permitted neurons to regenerate axons for 48hr. We analyzed Tuj1-stained axons and found that the average length of regrown axons was markedly increased in either the *Rtn4r* or *Atxn2* knockdown conditions relative to the non-targeting control; non-targeting-148.9µm, *Rtn4r*-188.8µm, and *Atxn2*-184.8µm (**Figure 4D and 4E**). These results provide evidence of a role for ataxin-2 in limiting regeneration after nerve injury and present ataxin-2 as a potential therapeutic target following nerve injury. Additionally, this finding provides further evidence that RNA-binding proteins that drive RNA granule formation following stress can limit axon regeneration— as has been reported for TIAR-2 (Andrusiak et al., 2019, Becker et al., 2017, Liu-Yesucevitz et al., 2011).

**Figure 4:**
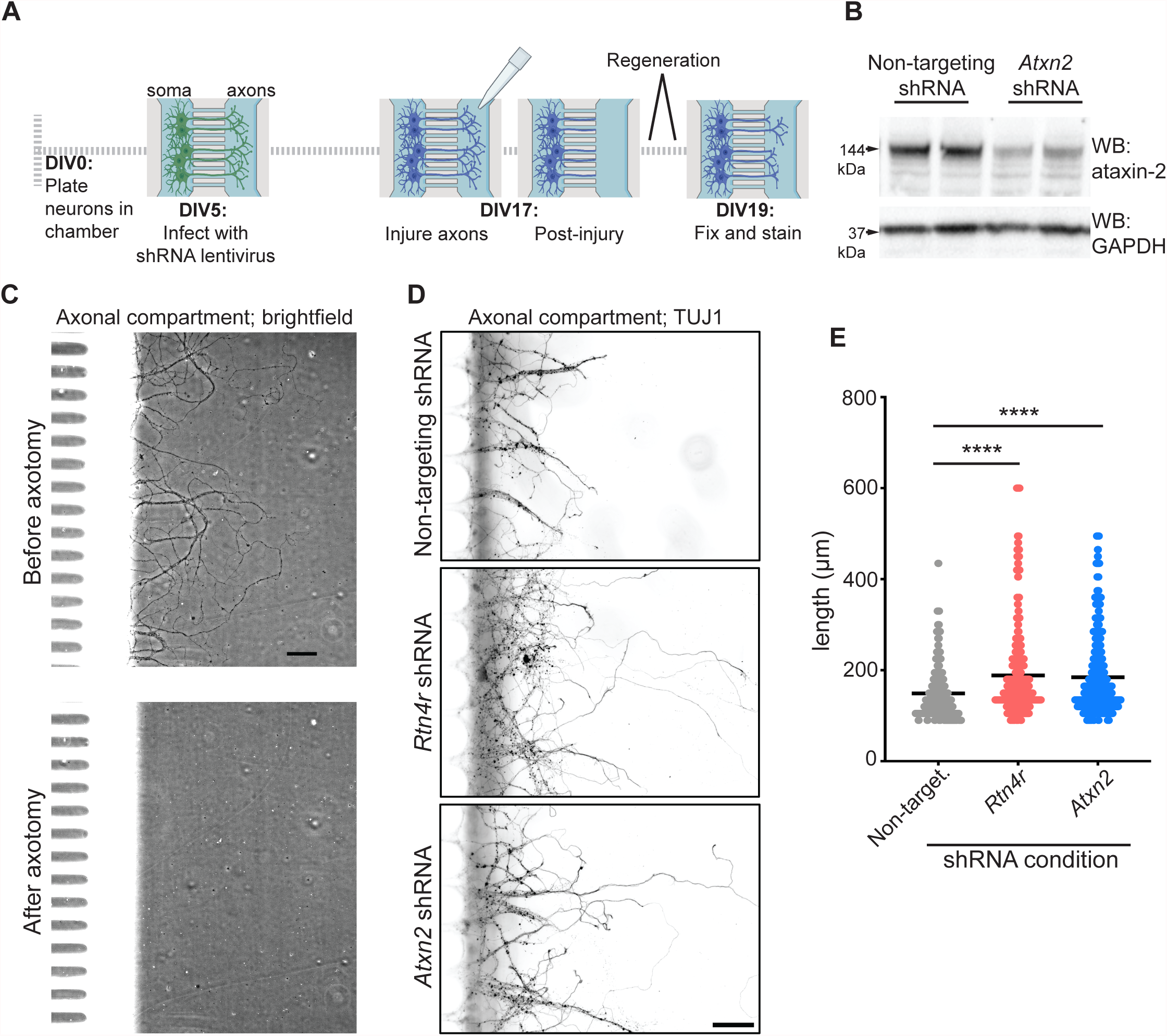
Reduction of ataxin-2 increases axonal regrowth after axotomy. **(A)** Timeline for axotomy and regeneration experiment using mouse primary neurons grown in microfluidics chambers. **(B)** Immunoblot on lysates from mouse neurons treated with *Atxn2* shRNA. **(C)** Brightfield images of an inner chamber of the axonal compartment of a microfluidics chamber before and after vacuum-assisted axotomy. Scale bar=50μm. **(D)** Immunocytochemistry and fluorescence microscopy on the axonal compartment after 48 hours of regrowth after axotomy. Tuj1 labels axons. Scale bar=50μm. **(E)** Quantification of the length of regenerating neurites (identified by the morphological presence of a growth cone) from three separate chambers per condition. One-way ANOVA with multiple comparisons. ****p ≤ 0.0001. Error bars represent ± SEM.

## Discussion

Here we performed an unbiased genome-wide screen and discovered RTN4/NoGo-Receptor as a novel regulator of ataxin-2 levels. We provide evidence that *RTN4R* functions upstream of ataxin-2. Since ataxin-2 has been implicated in two neurodegenerative diseases—ALS and SCA2— efforts are underway to target it therapeutically including the use of ASOs targeting *ATXN2* (Becker et al., 2017, Scoles et al., 2017). Together with the accompanying manuscript by Kim *et al*. identifying Etidronate, a potent small molecule inhibitor of ataxin-2 levels (please see accompanying manuscript), we now present an additional way to reduce ataxin-2 levels. RTN4/NoGo-Receptor seems like an optimal target for therapeutic reduction of ataxin-2 because in addition to gene-based strategies there are several ways in which it may be targeted, including receptor inhibition—demonstrated here—as well as neutralization of its ligands (Schwab, 2010). A RTN4/NoGo-Receptor Decoy (Wang et al., 2011, Wang et al., 2020) is being investigated in a clinical trial for chronic spinal cord injury (ClinicalTrials.gov identifier: NCT03989440). With additional targets and strategies to lower ataxin-2 levels in hand, combination therapies can be envisioned to have maximum therapeutic benefit and to mitigate potential negative effects of relying on a single target.

### Limitations of the study

The effectiveness of *RTN4R* knockdown in rescuing degeneration in a mouse model of ALS remains to be tested. However, reducing *ATXN2* is a validated method for rescuing degeneration phenotypes in a mouse model of TDP-43 toxicity and motor neuron degeneration (Becker et al., 2017) and in a mouse model of SCA2 (Scoles et al., 2017). Moreover, human genetics has provided compelling validation for *ATXN2* as a therapeutic target for both ALS and SCA2 (Elden et al., 2010; Scoles and Pulst, 2018). Future work will be needed to determine the precise method for reducing *RTN4R*—and subsequently *ATXN2*—in mouse models to test its influence on disease phenotypes. But given that clinical trials are underway to test a NoGo-Receptor inhibitor in spinal cord injury (ClinicalTrials.gov identifier: NCT03989440), lessons learned from that trial will hopefully aid in testing this approach for ALS and SCA2.

## Methods

### Cell culture and transfection

HEK293T cells were maintained in a 37°C incubator with 5% CO_2_ in DMEM with Glutamax and HEPES (Thermo Fisher Scientific, cat# 10564-029), 10% fetal bovine serum (vol/vol; Invitrogen, cat# 16000-044), and 1% Pen/Strep (vol/vol; Invitrogen, cat# 15140-122). Cells were reverse transfected on 96-well plates for luciferase assays. 25µL of Opti-MEM (Life Technologies, cat# 31985-062) with 0.12µL of Dharmafect 1 (Horizon Discovery, cat# T-2001-03) and 200nM (unless specified in figure) of ON-TARGETplus siRNA (Horizon Discovery) or non-targeting (Horizon Discovery, cat# D-001810-10) was added to an individual well and incubated at room temperature for 30 minutes. 1.0×10^4^ cells/well in 100µL of medium (without Pen/Strep) was added to the wells, and the plate was placed in the incubator for 72hr. To interrogate proteasome or autophagy regulation by HiBiT assay, 72hr after reverse transfection cells were treated for 6hr with 10µM MG-132 (Millipore Sigma, cat# M7449) or 24hr with 100µM Bafilomycin A (Sigma Aldrich, cat# B1793) or DMSO prior to lysing.

For immunoblotting, unedited HEK293T cells were maintained as above. Unedited cells were used to reduce cell line-specific effects. Cells were reverse transfected on 12-well plates. 200µL of Opti-MEM with 4µL of Dharmafect 1 and 200nM of ON-TARGETplus siRNA or non-targeting control was added to an individual well and incubated at room temperature for 30 minutes. 2.0×10^5^ cells/well were plated in 1mL of medium without Pen/Strep and placed in the incubator for 72hr prior to lysing for experimentation.

### Small-scale Luciferase Assays

Ataxin-2-HiBiT cells were lysed in-well with 125µL of Nano-Glo® HiBiT Lytic Buffer, and lysate from one well was split and used for detection of both HiBiT- and FFLuc-generated luminescence. 25µL of lysate was placed in two wells of an opaque white, flat bottom 96-well assay plate (Sigma Aldrich, cat# CLS3990). For HiBiT detection, 25µL of HiBiT lytic reagent (1:25 substrate, 1:50 LgBiT) was added to the lysate (Promega, cat# N3050). For FFluc detection, 25µL of ONE-Glo Assay Buffer (Promega, cat# E6120) was added to the lysate. Plates were incubated in the dark with gentle rotation for 10 minutes, then luminescence was measured on a Tecan Spark plate reader (Tecan). For all assays, HiBiT signal was normalized to the FFluc signal for each individual well of the original transfected plate then normalized to the non-targeting/untreated control, this value is represented in bar graphs.

### Genome Editing

HEK293T cells (ATCC) were transfected using Lipofectamine™ CRISPRMAX™ Cas9 Transfection Reagent (Thermo Fisher Scientific, cat# cmax00003), TrueCut Cas9 Protein V2 (Thermo Fisher Technology, cat# A36497), purified sgRNA and ssDNA (IDT). Sequences are listed in **Table 1**. 72hr later, cells were lifted and re-plated at a density of <1 cell/well in a 96-well plate. Wells were grown to confluency and split onto an additional plate. A HiBiT assay was performed on the additional plate as a preliminary screen for successful knock-in. The wells with HiBiT signal above the negative control (unedited cells) were maintained and confirmed for knock-in by Sanger sequencing (see **Table 1** for primers). For firefly luciferase integration, the successful clone was transduced with lentivirus using pLenti CMV V5-LUC Blast (Addgene, cat# w567-1), followed by blasticidin selection. Transduced cells were pooled and saved for future experimentation.

**Table 1:**
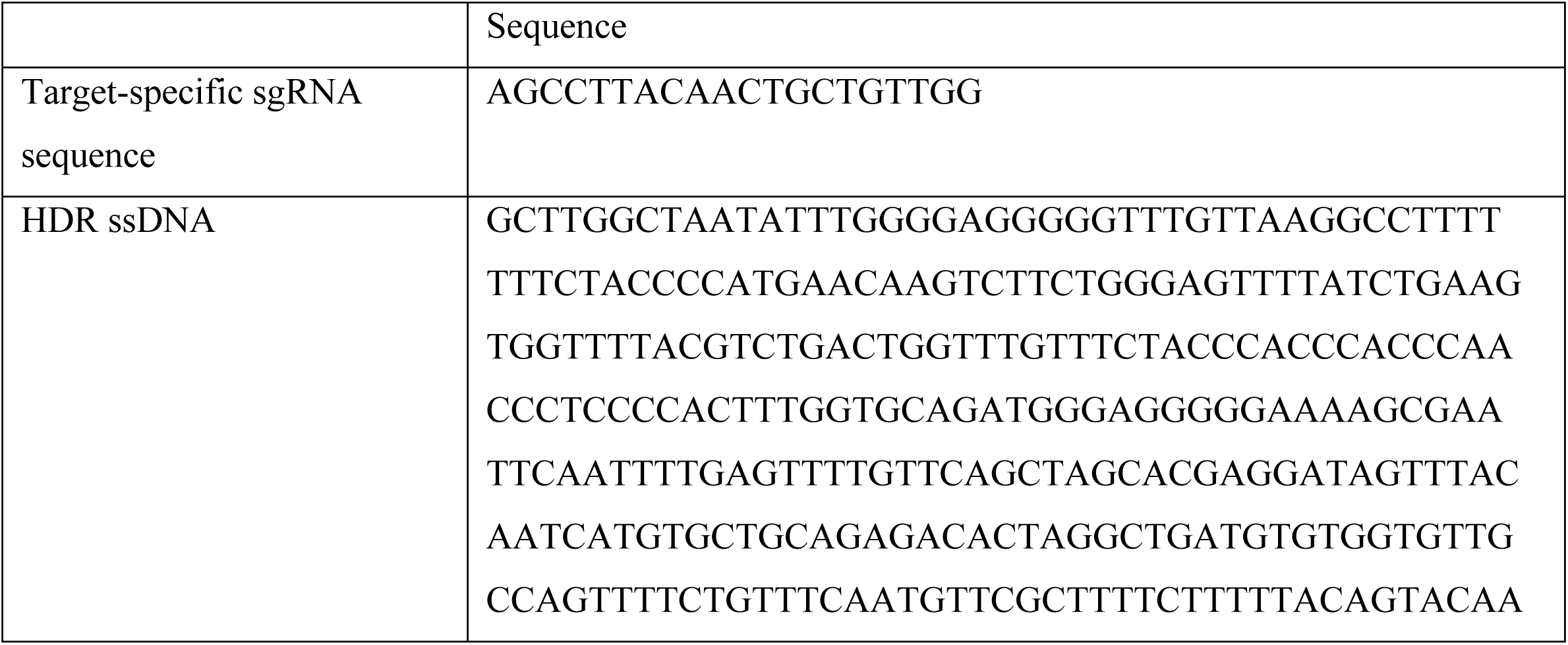
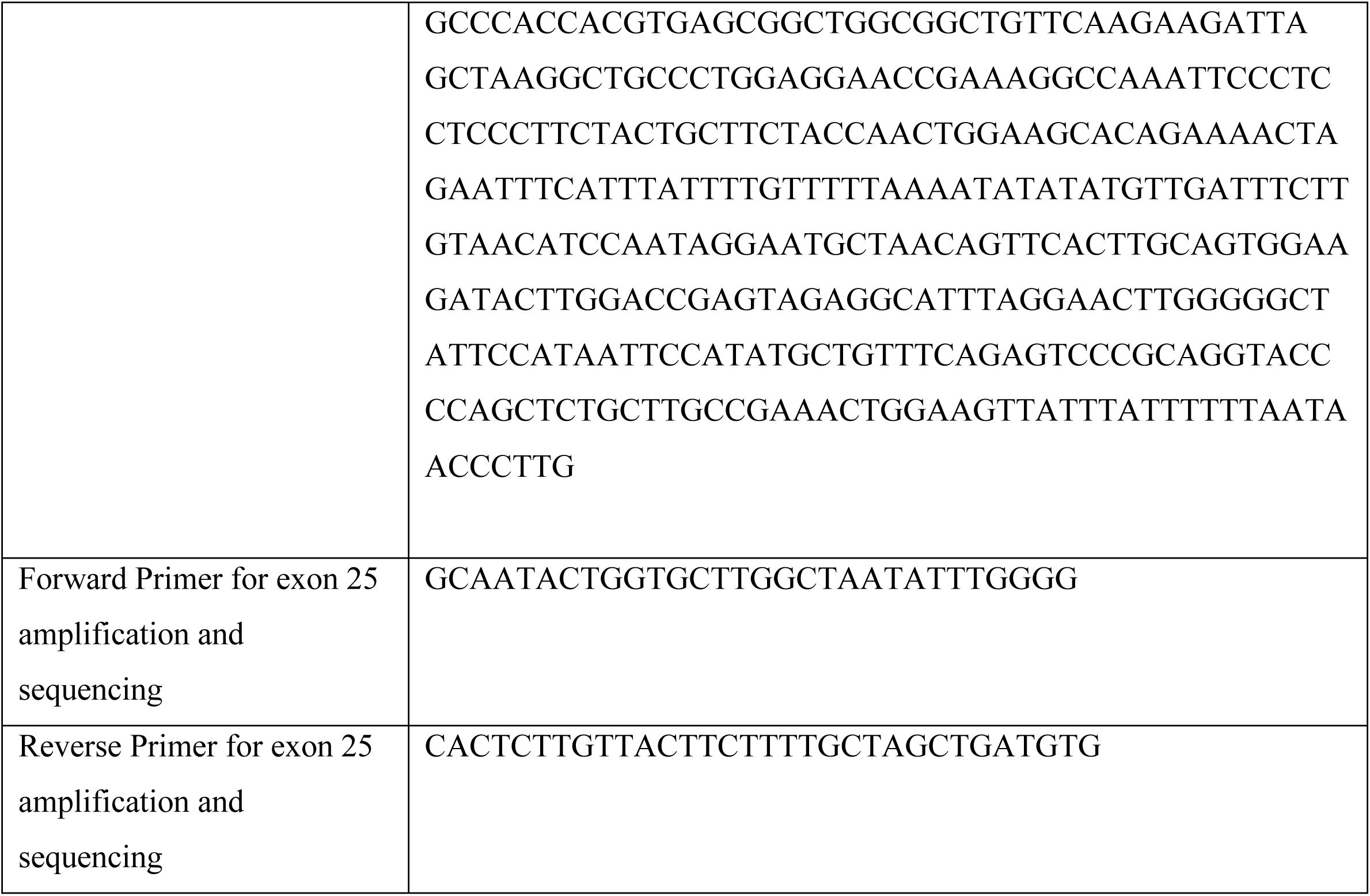

### Immunoblotting

Cells were lysed in RIPA buffer (Sigma Aldrich, cat# R0278) supplemented with Complete mini protease inhibitor tablet (EMD Millipore, cat# 11836170001) and clarified by centrifugation at 21,000 x *g* for 15 minutes. Protein concentration was measured by BCA protein assay (Thermo Fisher Scientific, cat# 23225). Lysates were diluted to equal protein concentration in 1X NuPAGE® LDS Sample Buffer (Life Technologies, cat# NP0008). Lysates were boiled at 70°C for 10 minutes, then loaded on a NuPAGE® Novex® 4-12% Bis-Tris Protein gel (Life Technologies, cat# NP0321). Protein was transferred onto a nitrocellulose membrane (Bio-Rad, cat# 162-0115) at 4°C for 1hr and 45 minutes in 1X NuPAGE® Transfer Buffer (Life Technologies, cat# NP0006-1). Blocking and antibody incubation was performed in 2% BSA (Sigma Aldrich, cat# A7906) in PBS + 0.1% tween-20 (PBST). Washes were performed in PSBT. Primary antibodies were used at 1:1000, this includes: ataxin-2 antibody (Novus, cat# NBP1-90063), Tuj1 antibody (BioLegend, cat# 802001), actin antibody (EMD Millipore, cat# MAB1501), RTN4/NoGo-Receptor antibody (Abcam, ab184556), and GAPDH antibody (Sigma Aldrich, cat# G8795). Goat anti-Rabbit HRP (Life Technologies, cat# 31462) or goat anti-mouse HRP (Thermo Fisher Scientific, cat# 62-6520) secondary antibodies were used at 1:5000. Membranes were developed in ECL Prime (Sigma Aldrich, cat# GERPN2232), and imaged on a Bio-Rad ChemiDoc XRS+ imager (BioRad). HiBiT-based immunoblotting was performed according to the Nano-Glo® HiBiT Blotting System protocol (Promega, cat# n2410).

For mouse cortex, male and female mice at P20 were sacrificed for tissue harvesting. Ice cold RIPA buffer with protease inhibitor was added and tissue was homogenized using a motorized pestle. Crude lysates were rocked at 4°C for 30 minutes, passed through a homogenization column (Qiagen, cat# 79656), and centrifuged at 21,000 x *g* for 15 minutes. Protein was quantified in the clarified supernatant and prepared for immunoblotting as above.

### Whole-genome siRNA screen

The whole-genome siRNA screen was performed in the High-Throughput Bioscience Center (HTBC) at Stanford University. Briefly, we used the Dharmacon Human Genome siARRAY library (cat# G-005000-025) to target 21,121 genes individually in a single well of a 384-well plate with 4 pooled siRNAs. All siRNA pools were tested in duplicate for both FFluc and HiBiT on 4 separate plates. Each plate had three controls: siTOX control siRNA, nontargeting control siRNA, and *ATXN2* siRNA (Horizon Discovery, cat# L-011772-00) as a toxicity, negative, and positive control, respectively. All siRNAs were reverse transfected using 10µL of Opti-MEM, 0.075µL/well of Dharmafect 1, and 10µL siRNA (final concentration of 50nM). Cells were seeded at 1000 cells/well in 30µL of media (as above without Pen/Strep) in solid white 384-well plates, then placed in a 37°C incubator with 5% CO_2_. After 3 days, plates were removed from the incubator. HiBiT lytic reagent was made by diluting Nano-Glo® HiBiT Lytic Substrate (1:50) and LgBiT Protein (1:100) in Nano-Glo® HiBiT Lytic Buffer. For HiBiT detection, 10 µL of HiBiT lytic reagent was dispensed into wells with a Multidrop Reagent Dispenser (Thermo Scientific), and rapidly shook for 15 seconds. For FFluc detection, 10µL of ONE-Glo Assay Buffer was dispensed into wells and rapidly shook for 15 seconds. Luminescence was read on a Tecan Infinite M1000 Pro (Tecan).

Z-scores were calculated for each siRNA replicate using the in-plate standard deviation, then averaged for both HiBiT and FFluc conditions respectively. The average Z-scores were used to filter for primary screen hits. siRNAs were only considered if they had an average FFluc Z-score greater than −1. siRNAs were categorized as negative regulators of ataxin-2 expression if they had an average HiBiT Z-score less than −1.65. 102 of these were retested in a secondary screen by handpicking siRNAs from the original library stock. The same experimental parameters applied, except for the performance of a single replicate for FFluc detection. High confidence hits were chosen if they had a greater than 30% decrease in average HiBiT levels relative to the non-targeting siRNA control run in parallel on the same plate. High confidence hits were ranked by average HiBiT Z-score, then were filtered further to determine the most optimal therapeutic target. These were filtered out if they had an average gene effect less than −1 as determined by DEMETR2 or CERES (depmap.org) or GTEx cortical pTPM less than 15 (proteinatlas.org).

### RNA Quantification

Cells were reverse transfected with siRNA in 12-well plates as described above. RNA was isolated using the PureLink® RNA Mini Kit according to the kit protocol with DNase digestion (Thermo Fisher Scientific, cat# 12183025). 500ng of RNA was used to make cDNA using the Applied Biosystems High-Capacity cDNA Reverse Transcription kit (Thermo Fisher Scientific, cat# 4368813). qPCR was performed in a 20µL reaction using TaqMan™ Universal Master Mix II (Thermo Fisher Scientific, cat# 4440040) and 25ng of RNA. 1µL of 20X TaqMan gene-specific expression assay was added to the reaction (Thermo Fisher Scientific; human *ATXN2*: Hs00268077_m1, human *ACTB:* Hs01060665_g1, human *RTN4R*: Hs00368533_m1, mouse *ATXN2*: Mm00485932_m1, mouse *GAPDH*: Mm99999915_g1, mouse *RTN4R*: Mm00452228_m1). All conditions were run in technical triplicates that were averaged to account for each of the biological triplicates. Thermocycler was programmed according to the suggested protocol. Relative expression was calculated using the ΔΔCt method.

### Lentivirus production

HEK293T cells were grown on 10cm culture dishes to 80-90% confluency. Cells were transfected using Lipofectamine 3000. Briefly, 10µg of shRNA vector (Mission® shRNA, Sigma Aldrich), 2.9µg pRSV-REV, 5.8µg pMDLg/pRRE, 3.5µg pMD2.G, and 40µL of P3000 was added to 1mL of Opti-MEM. This was combined with another 1mL of Opti-MEM with 40µL of Lipofectamine 3000. This mixture was incubated at room temperature for 10 minutes then added to cells in media without antibiotics. 48hr after transfection, media was removed from cells, passed through a 0.45µm PES filter, and combined with Lenti-X concentrator (Takara, cat# 631232). The mixture was incubated over night at 4°C. This was centrifuged at 1,500 x *g* for 45 minutes at 4°C. The pellet was resuspended in 1mL Neurobasal Medium (Life Technologies, cat # 21103-049), 50µL was added directly to cells for transduction.

### Primary neuron culture

Animals were bred, cared for, and experimented on as approved by the Administrative Panel of Laboratory Animal Care (APLAC) of Stanford University. Time-pregnant C57BL/6 mice were procured from Charles River Labs. Mouse embryos were removed at E16.5, and cortices were isolated and placed in ice cold Dissociation Media (DM; PBS without Mg^2+^ or Ca^2+^, supplemented with 0.6% glucose, 10mM HEPES, and Pen/Strep). Neurons were dissociated with the Papain Dissociation System (Worthington Biochemical Corporation, cat# LK003153). Cells were seeded on Poly-L-lysine (Sigma Aldrich, cat# P4832) coated 24-well plates at a density of 350,000cells/well. They were grown in Neurobasal media supplemented with B-27 at 1:50 (Life Technologies, cat# 17504-044), Glutamax at 1:100 (Invitrogen, cat# 35050-061) and Pen/Strep. Neurons were maintained in a 37°C incubator with 5% CO_2_ with half media changes every 4 to 5 days.

At DIV 5, neurons were transduced with virus as above (Mission® shRNA, mouse *RTN4R*: TRCN0000436683). 24hr later, a half media change was performed. Neurons were maintained for 12 days after transduction prior to harvesting for experimentation. For NEP1-40 (Tocris, cat# 1984) treatment, 1mg of NEP1-40 was diluted in 21.62µL of DMSO and 843.2µL of nuclease-free water (Thermo Fisher Scientific, cat# AM9937) to a working concentration of 250µM. The peptide was added to the final concentration specified along with vehicle (0µM). Cells were maintained for 48hr prior to experimentation.

For microfluidics chamber experiments, primary neurons were isolated as above and plated on one side of a Poly-D-lysine coated microfluidics chamber (Xona Microfluidics, cat# XC450) with a microchannel length of 450µm at a density of 200,000cells/chamber. Media was changed the next day, then half-changed every 3 days following. Lentivirus was added to the somatic compartment as above, then maintained for 12 days prior to axotomy. Vacuum-assisted axotomy was achieved by complete aspiration of media from the axonal compartment/inner chamber, allowing for an air bubble to dislodge and shear axons(Tong et al., 2015). Media was replaced and completely aspirated a second time. Media was replaced carefully to prevent the inclusion of any air bubbles in the inner chamber. A half media change was performed the next day. Neurons were allowed to regrow axons for a total of 48hr prior to fixation.

### iNeuron culture

We induced neuron differentiation in human iPSC-derived neurons (iNeurons) using a previously established protocol with a Tet-On induction system that allows expression of the transcription factor NGN2(Bieri et al., 2019). Briefly, iPS cells were maintained with mTeSR1 medium (Stemcell Technologies, cat# 85850) on Matrigel (Fisher Scientific, cat# CB-40230). On the next day of passage, NGN2 was expressed by adding doxycycline (2µg/ml) and selection with puromycin (2µg/ml) for rapid and highly efficient iNeuron induction. Three days after induction, iNeurons were dissociated and grown in Neurobasal medium containing N-2 supplement (Gibco, cat# 17502048), B-27 supplement, BDNF (R&D Systems) and GDNF (R&D Systems) on Matrigel-coated plates.

At 7 days post neural induction, iNeurons were transduced with virus as above (Mission® shRNA, human *RTN4R*: TRCN0000061558). 24hr later, a half media change was performed. iNeurons were maintained for 3 days before being re-transduced with a half dose of virus, then maintained for another 5 days prior to harvesting for experimentation. For NEP1-40 treatment, 1mg of NEP1-40 was diluted as above. The peptide was added at 7 days post induction to the final concentration specified along with vehicle (0µM). Cells were maintained for 48hr prior to lysis.

### Immunocytochemistry and fluorescence microscopy

Cells were fixed for 10 minutes in a solution of 4% paraformaldehyde/4% sucrose in PBS-MC (PBS with 1mM MgCl_2_ and 0.1 mM CaCl_2_). Then cells were washed in PBS-MC, and permeabilized for 10 minutes in 0.1% Triton-X in PBS-MC. Cells were blocked in 2% BSA for an hour, then incubated in primary antibody for at least 2 hours at room temperature [ataxin-2 (Novus), MAP2 (Synaptic Systems, cat# 188004), Tuj1 (BioLegend, cat# 802001), NeuN (EMD Millipore, cat# MAB377)]. Cells were washed 3 times with PBS-MC, followed by incubation in species-specific Alexa Fluor®-labeled secondary antibody (Thermo Fisher Scientific) for 1 hour at 1:1000. Cells were subsequently washed and placed in ProLong Gold Antifade with DAPI (Thermo Fisher Scientific, cat# P36931), and imaged using a Leica DMI6000B fluorescent microscope.

Images were processed in ImageJ. For neuronal expression, individual MAP2 labeled soma were selected as regions of interest (ROIs), in which ataxin-2 fluorescence was measured and averaged within each frame. This was performed for 4 frames per biological replicate (4 replicates for the NEP1-40 treatment and 3 for the vehicle), totaling 16 frames for the NEP1-40 condition and 12 for the vehicle. For stress granule analysis, stress granules were automatically determined based on shape and size using the Analyze Particles function in ImageJ in the G3BP channel. Each stress granule was made into an ROI in which TDP-43 fluorescence was measured and averaged for every stress granule in the frame. This was performed for 6 frames per biological duplicate, totaling 12 frames per condition. For axonal regrowth after axotomy, entire axonal compartments were imaged for three microfluidics chambers per shRNA condition (9 total). A grid was superimposed onto each image with demarcations every 15µm after the end of the microchannels. The length of every Tuj1-labeled neurite with a growth cone was determined by their crossing point on the grid and binned into lengths. The average of all neurites from each condition was taken.

### Statistical analyses

Statistical analyses were performed in GraphPad Prism v.9, except the screen data calculations, which was performed in MATLAB (MathWorks). An unpaired Student’s t-test (two tailed) with a 95% confidence interval (CI) was performed for all assays comparing two experimental groups. A two-way analysis of variance (ANOVA) with a 95% CI was performed for the HiBiT assays with multiple siRNA conditions compared to respective non-targeting controls run in parallel and siRNA/drug treatment. A one-way ANOVA with multiple comparisons was performed for all other experiments with more than two experimental conditions. A Fisher’s LSD test was performed for the NEP1-40 treated HiBiT assay. A Dunnett’s multiple comparisons test was performed for all others. All bar graphs show the mean ± SEM. All samples were randomly assigned to experimental groups. No statistical methods were used to predetermine sample sizes.

## Supplemental information

**Table S1:**
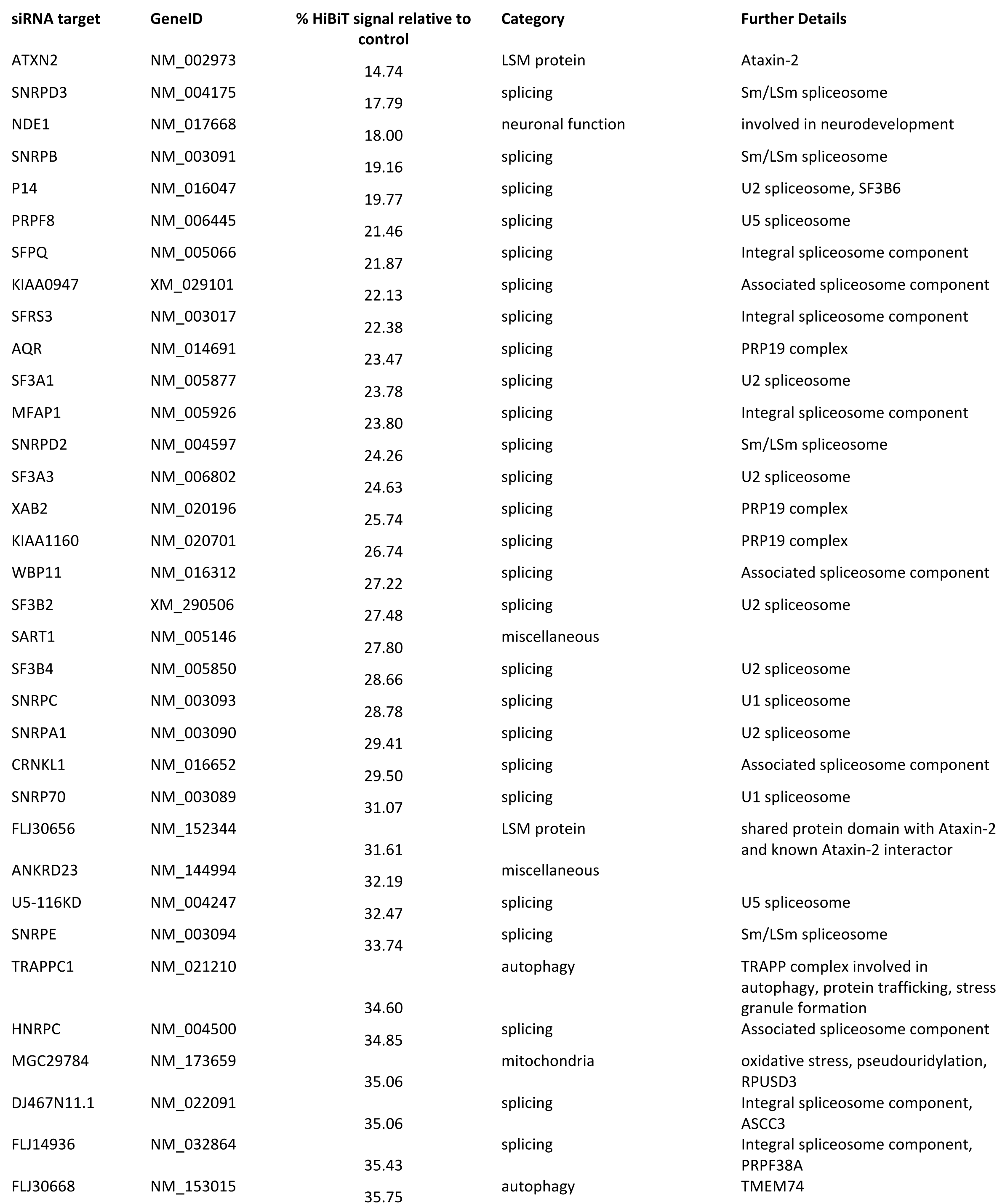
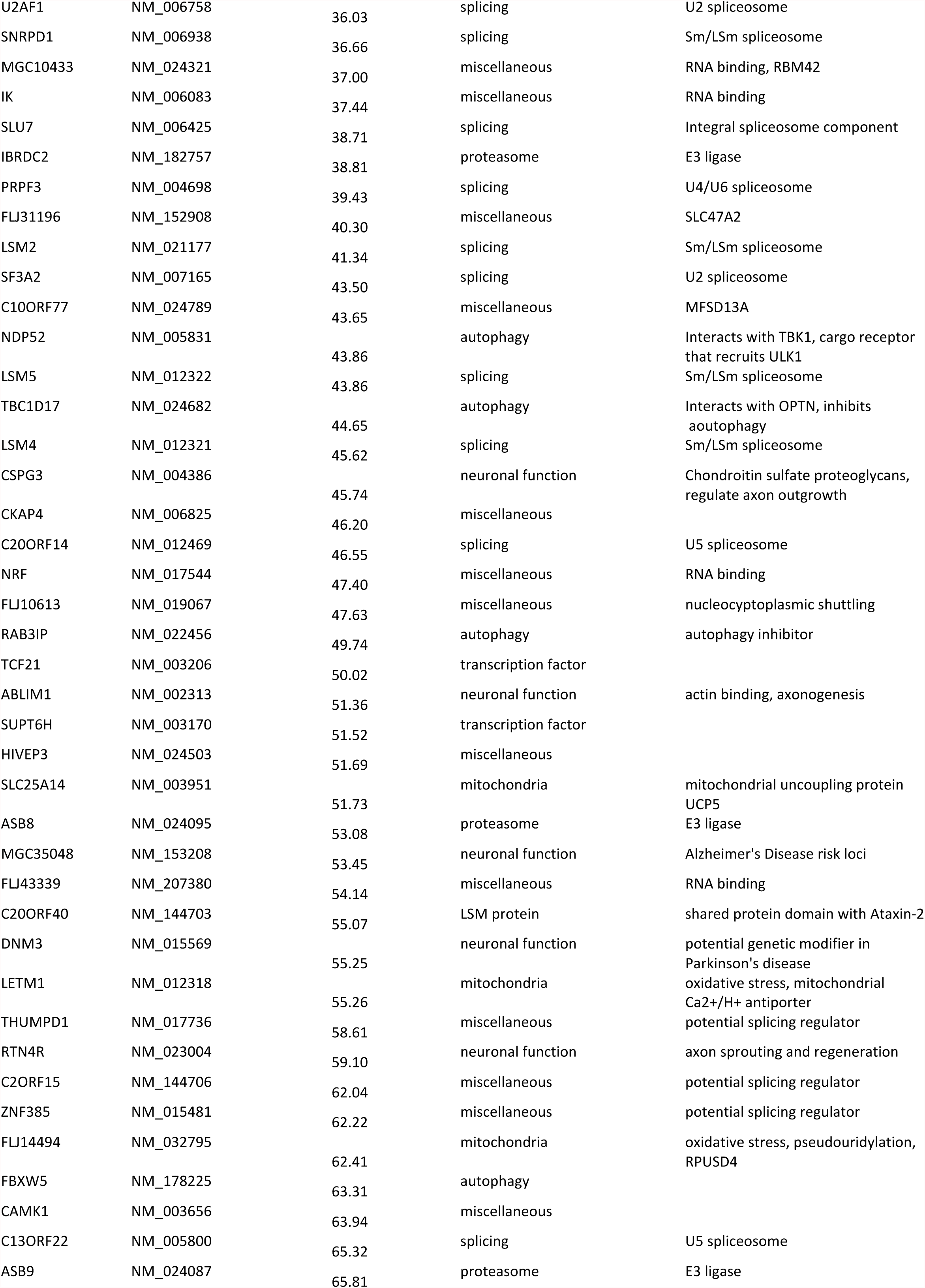
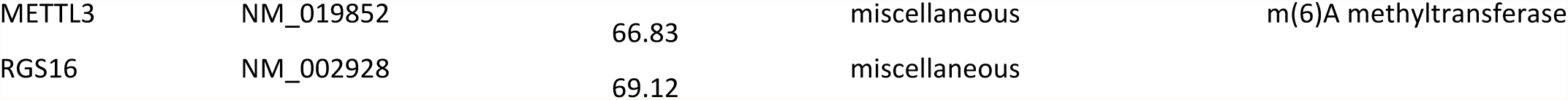
List of High Confidence Hits. % HiBiT signal relative to control was calculated from data collected in the secondary screen. Two biological replicates were performed for HiBiT signal and averaged and then normalized to the average signal from the non-targeting control wells run in parallel.

**Figure S1:**
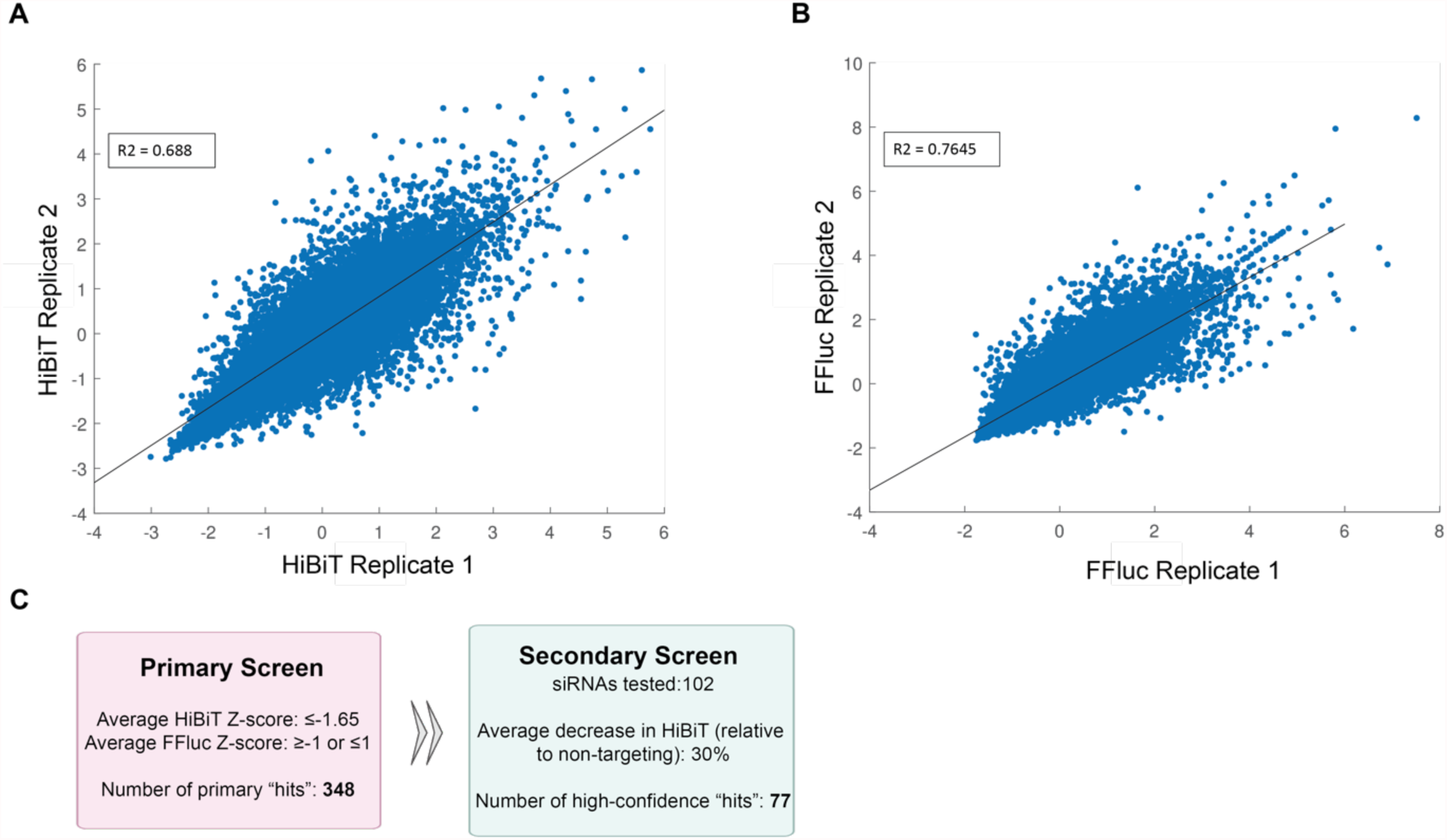
Ataxin-2 screen results and filters. Plots of the primary screen HiBiT data **(A)** and FFLuc data **(B)** for both replicates. R^2^= 0.688 and 0.7645, respectively. **(C)** Filters applied to screen results to determine the highest confidence ataxin-2 regulators.

**Figure S2:**
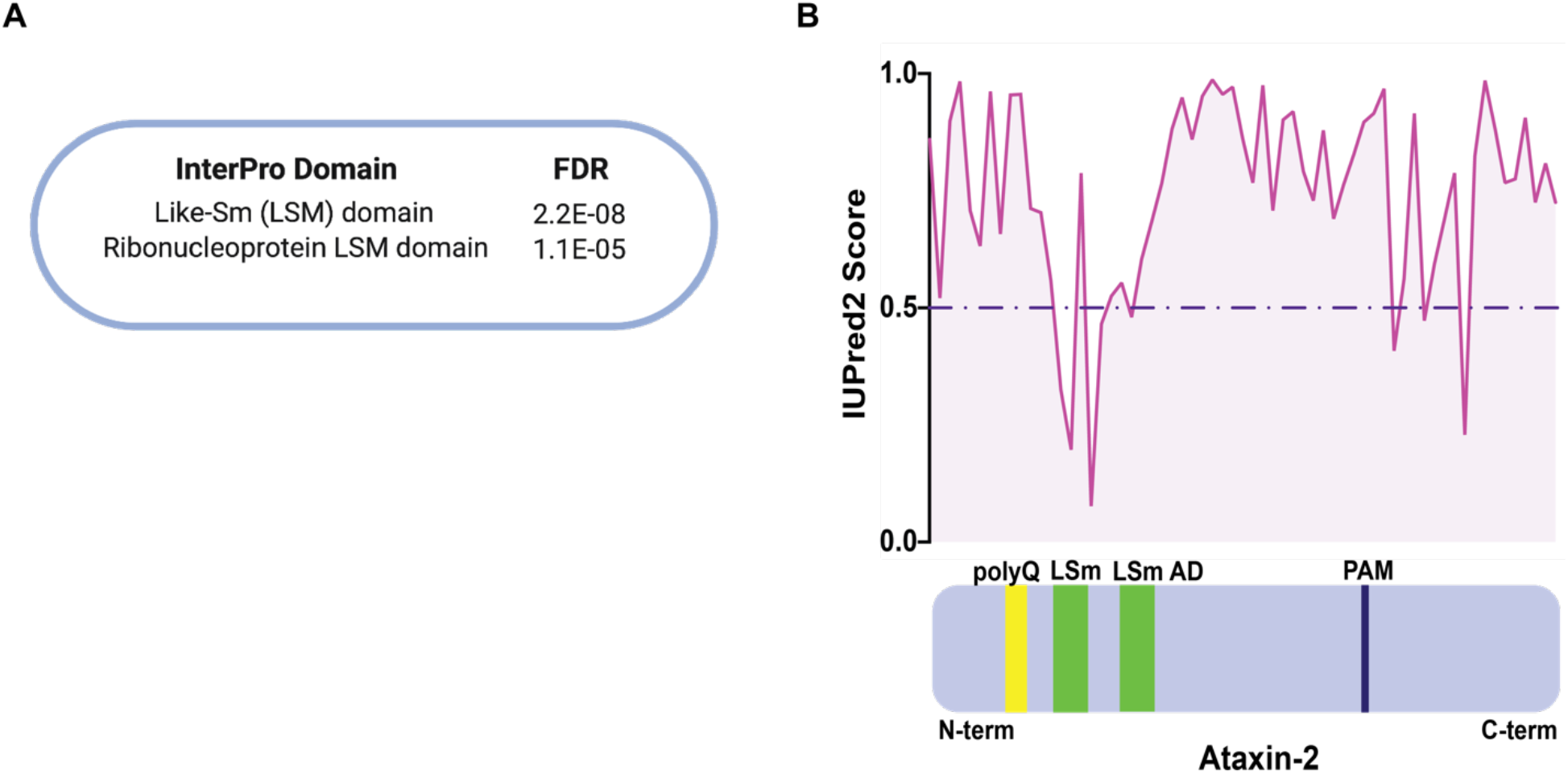
Shared protein domain between ataxin-2 and its regulators. **(A)** InterPro domain enrichment of primary screen hits. **(B)** Bottom: representation of ataxin-2 protein domains including the poly-glutamine stretch (polyQ), the LSm and LSm-associated (LSm AD) domains, and the PABP-interacting motif (PAM). Top: graph of the IUPred2 score, a prediction of protein disorder, for the amino acid sequence of ataxin-2.

**Figure S3:**
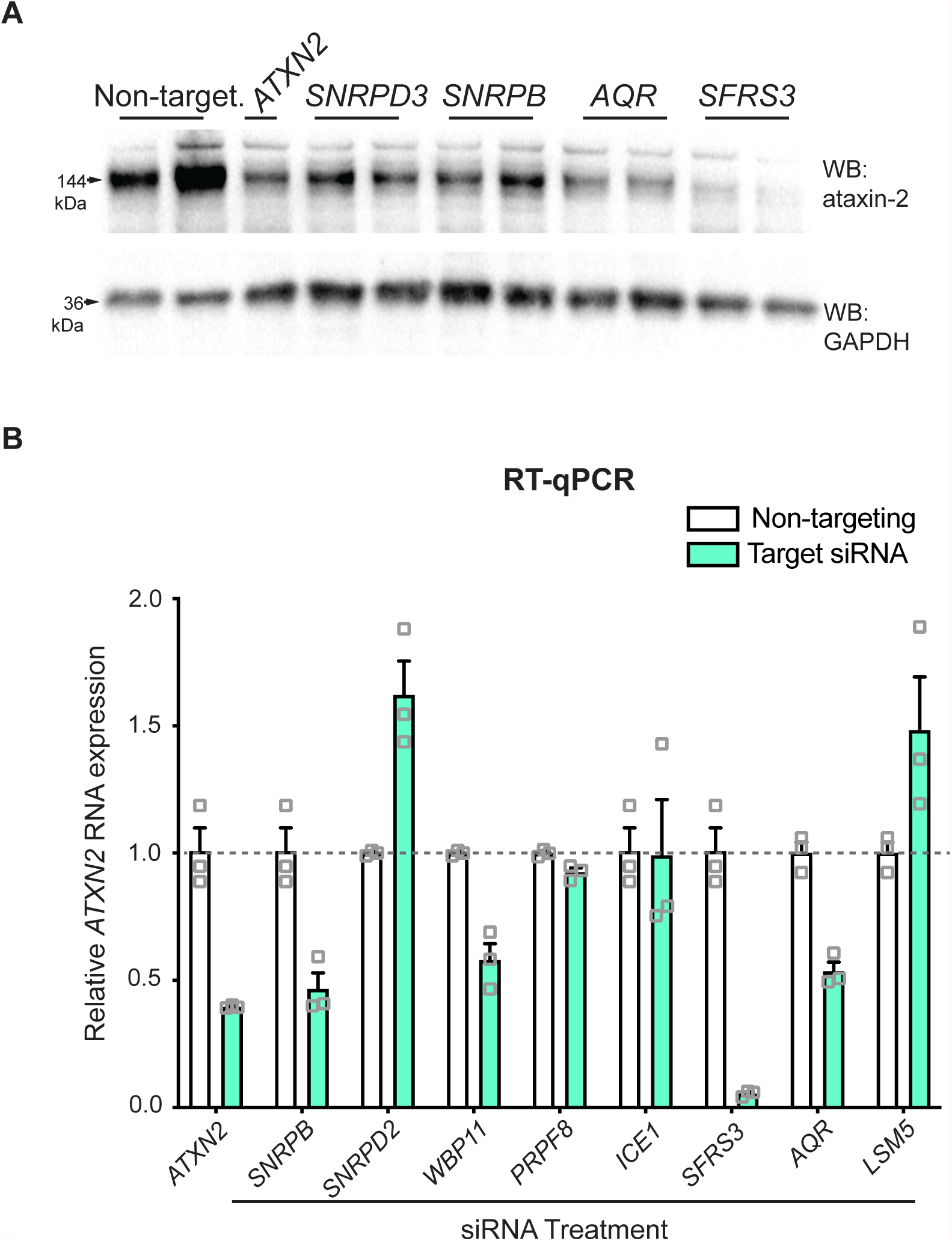
Further validation of screen results. **(A)** Immunoblot of ataxin-2 and GAPDH levels after siRNA treatment in unedited HEK293T cell lysates. **(B)** RT-qPCR on RNA from cells treated with non-targeting siRNA or siRNA targeting various splicing factors. We probed for *ATXN2* transcript along with *ACTB* for normalization. Two-way ANOVA with multiple comparisons: *WBP11* and *AQR*, p ≤ 0.05. *LSM5* and *SNRPB*, p ≤ 0.01. *ATXN2* and *SNRPD2*, p ≤ 0.001. *SFRS3*, p ≤ 0.0001. Error bars represent ± SEM.

**Figure S4:**
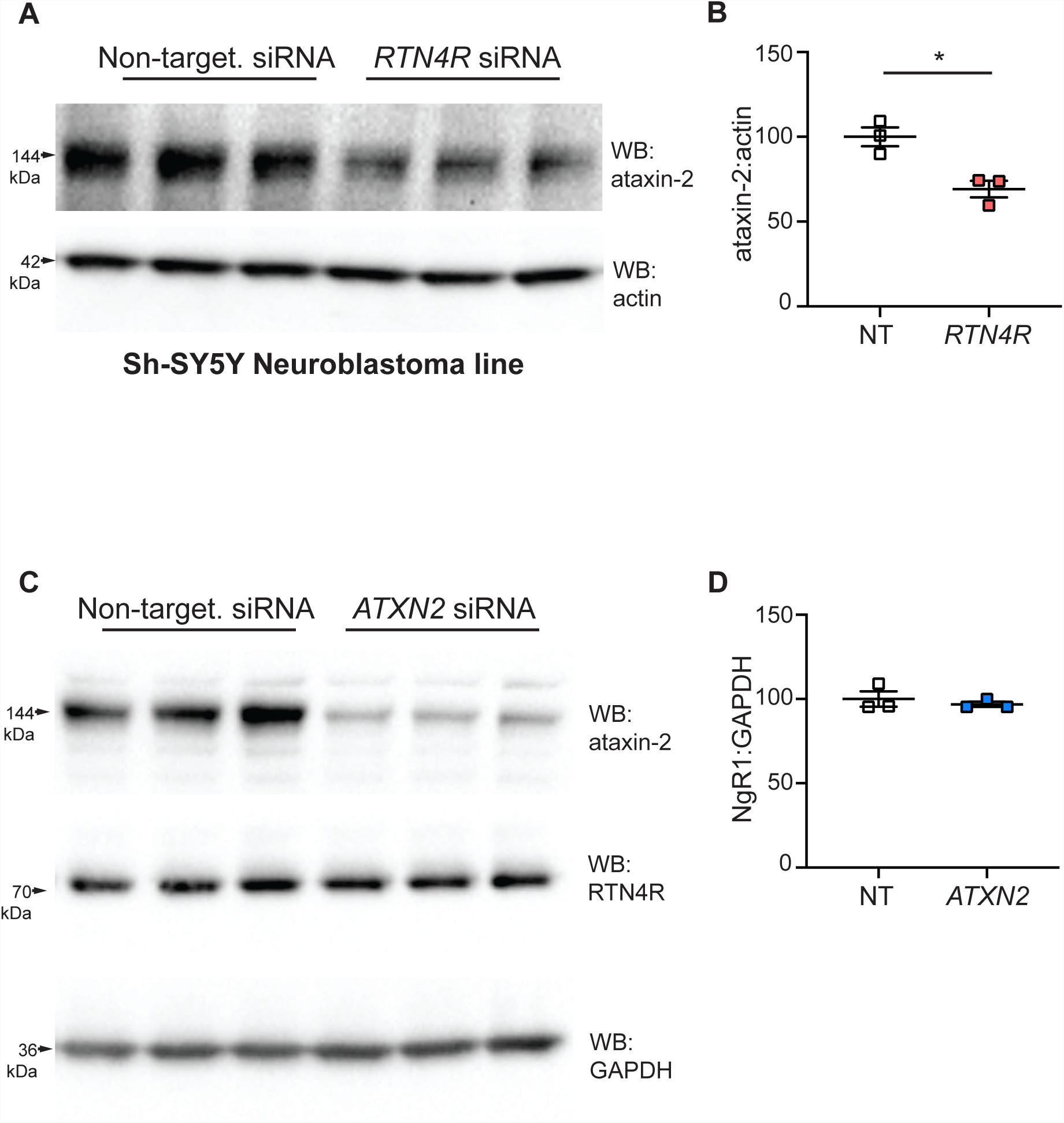
Further validation of *RTN4R* as a regulator of ataxin-2. **(A)** Immunoblot on SH-SY5Y cell lysates after *RTN4R* siRNA treatment. Ataxin-2 levels are quantified in **(B). (C)** Immunoblot on HEK293T cell lysates after *ATXN2* siRNA treatment. RTN4/NoGo-Receptor levels are quantified in **(D)**. Student’s t-test. *p ≤ 0.05. Error bars represent ± SEM.

**Figure S5:**
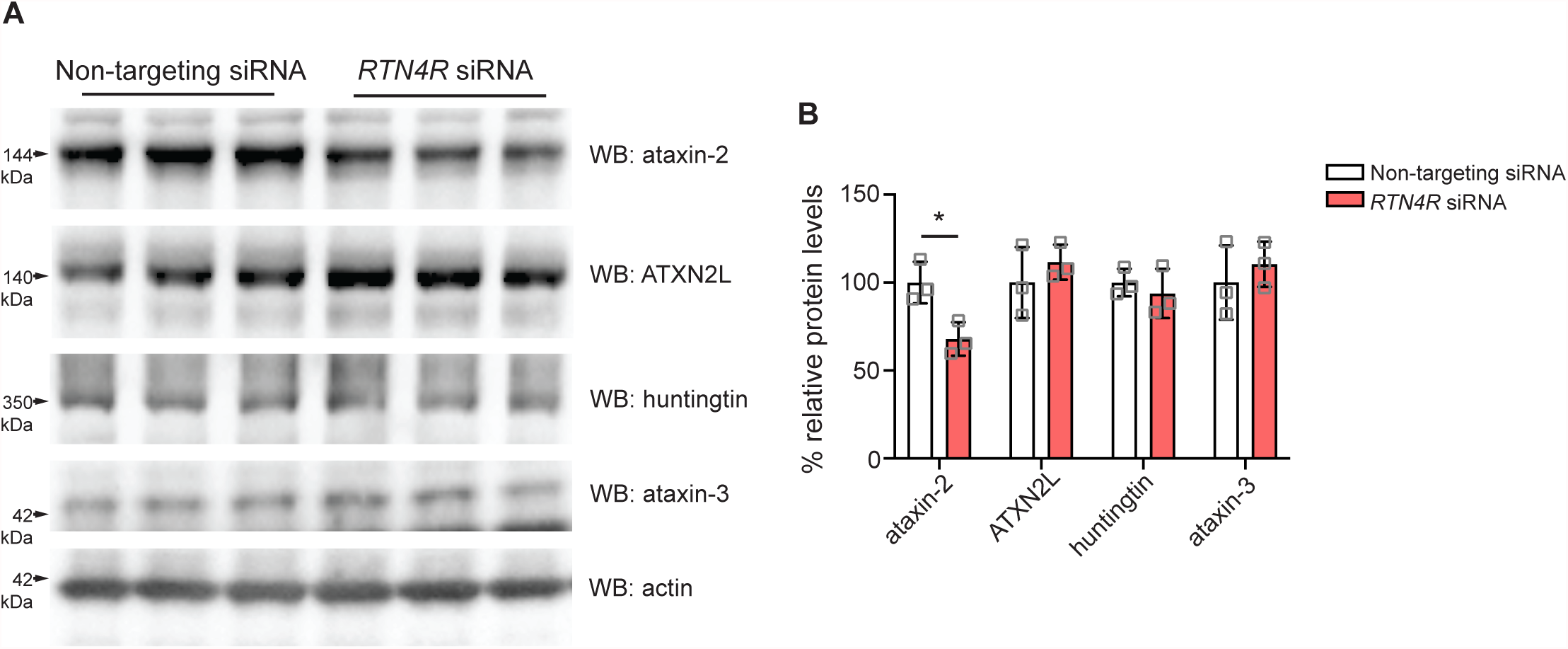
Knockdown of RTN4R does not alter the expression of other polyQ proteins or ATXN2L. **(A)** Immunoblot of ATXN2L (Ataxin-2 paralog) and polyQ disease proteins ataxin-2, huntingtin, and ataxin-3 after *RTN4R* siRNA treatment in unedited HEK293T cell lysates. Quantified in **(B)**. Student’s t-test. *p ≤ 0.05. Error bars represent ± SEM.

**Figure S6:**
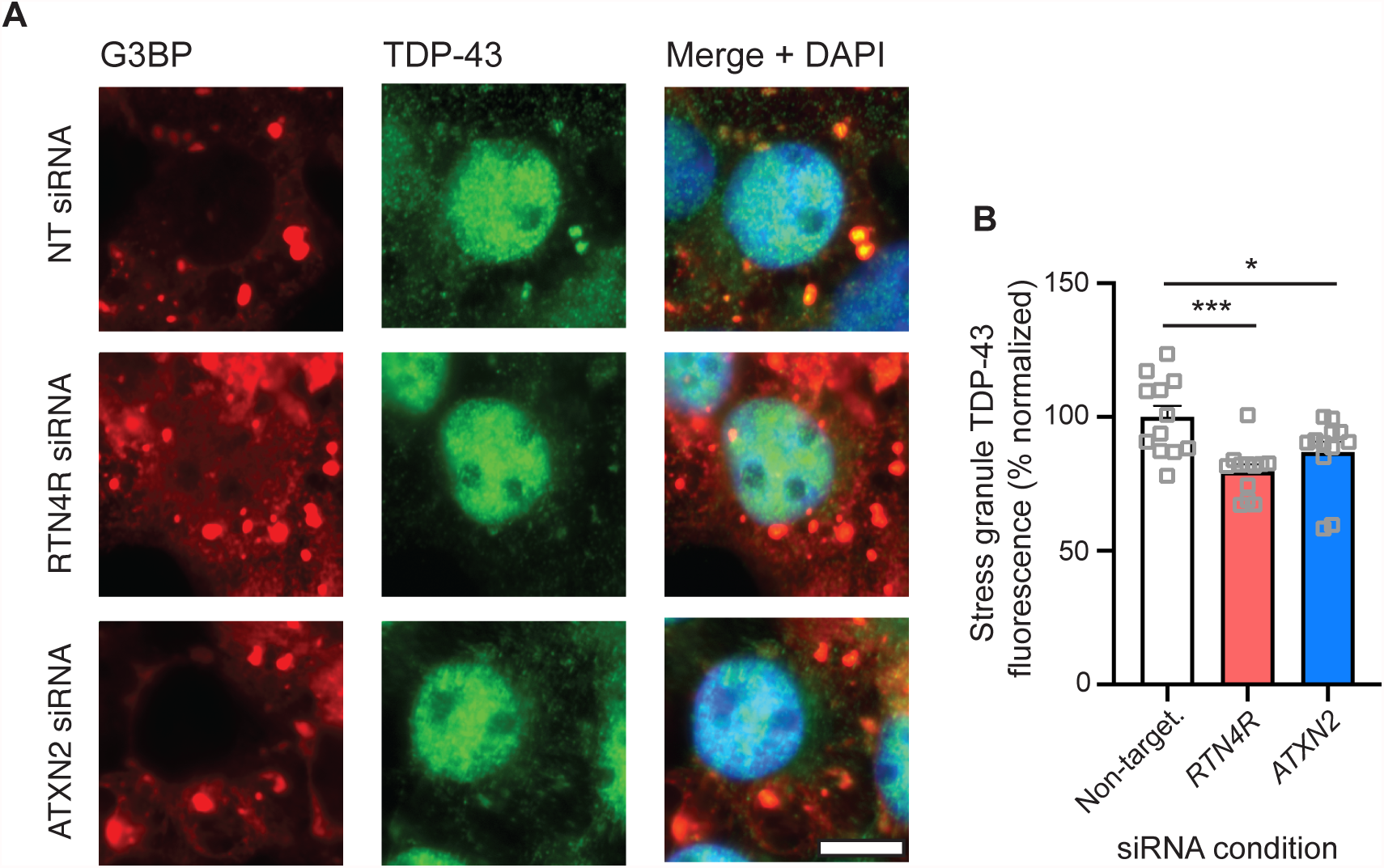
Knockdown of *RTN4R* reduces TDP-43 localization to stress granules. **(A)** We treated HEK293T cells with siRNA, and subsequently treated for 30 minutes with 0.5mM sodium arsenite to induce stress granules. We probed for TDP-43 and the stress granule marker G3BP via immunocytochemistry. Scale bar=10μm. The average fluorescence of TDP-43 in stress granules is quantified in **(B)**. One-way ANOVA, *p ≤ 0.05, ***p ≤ 0.001. Error bars represent ± SEM.

**Figure S7:**
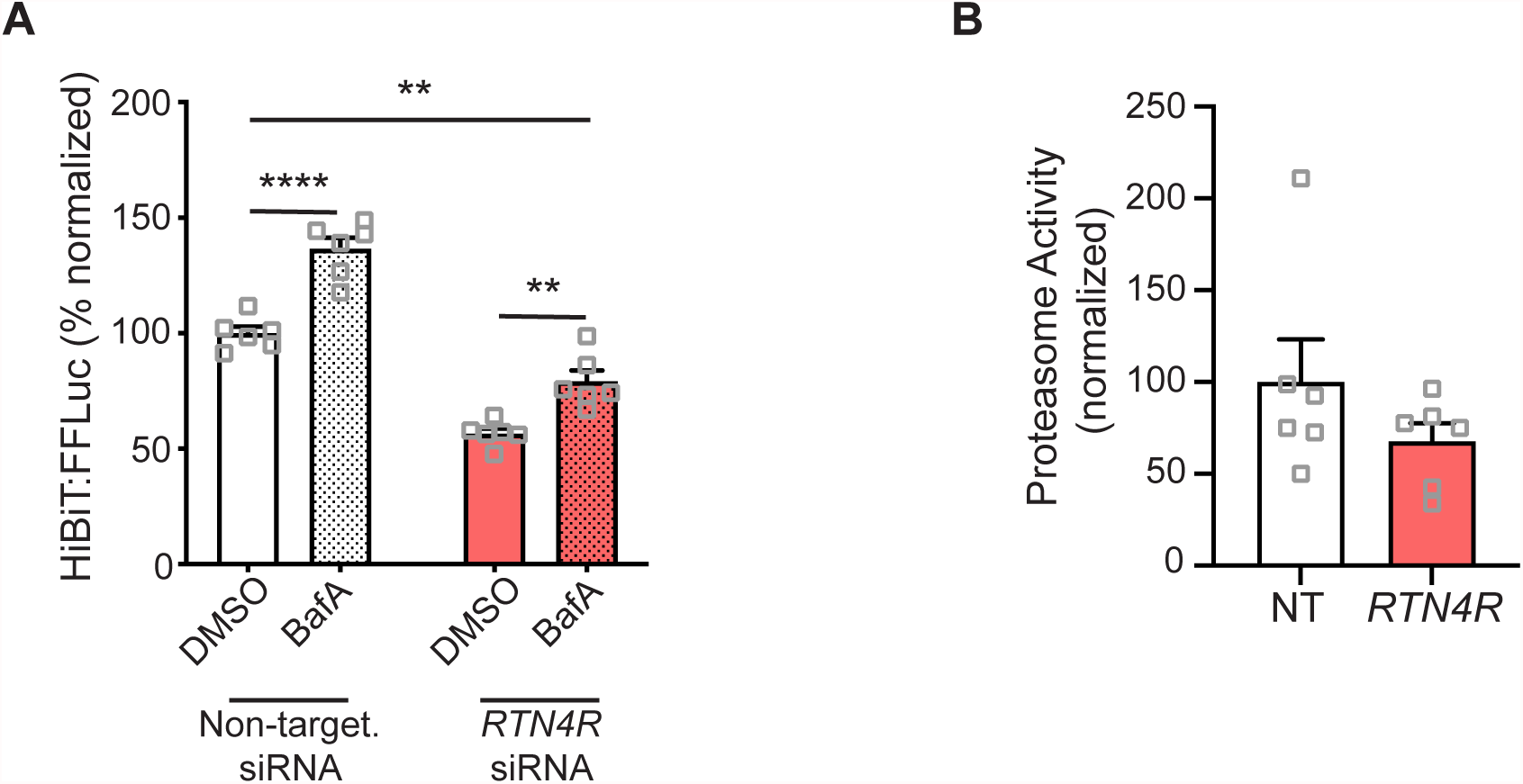
Knockdown of *RTN4R* does not decrease ataxin-2 through autophagy pathways and does not increase general proteasome activity. **(A)** We treated ataxin-2-HiBiT cells with siRNA, then treated for 24hr with autophagy inhibitor Bafilomycin A1 or DMSO. We performed a luciferase assay to measure HiBiT activity. **(B)** Proteasome activity assay (utilizing Suc-LLVY-AMC as a substrate) in HEK293T cell lysates after *RTN4R* siRNA treatment. Two-way ANOVA with multiple comparisons. **p ≤ 0.01, ****p ≤ 0.0001. Error bars represent ± SEM.

**Figure S8:**
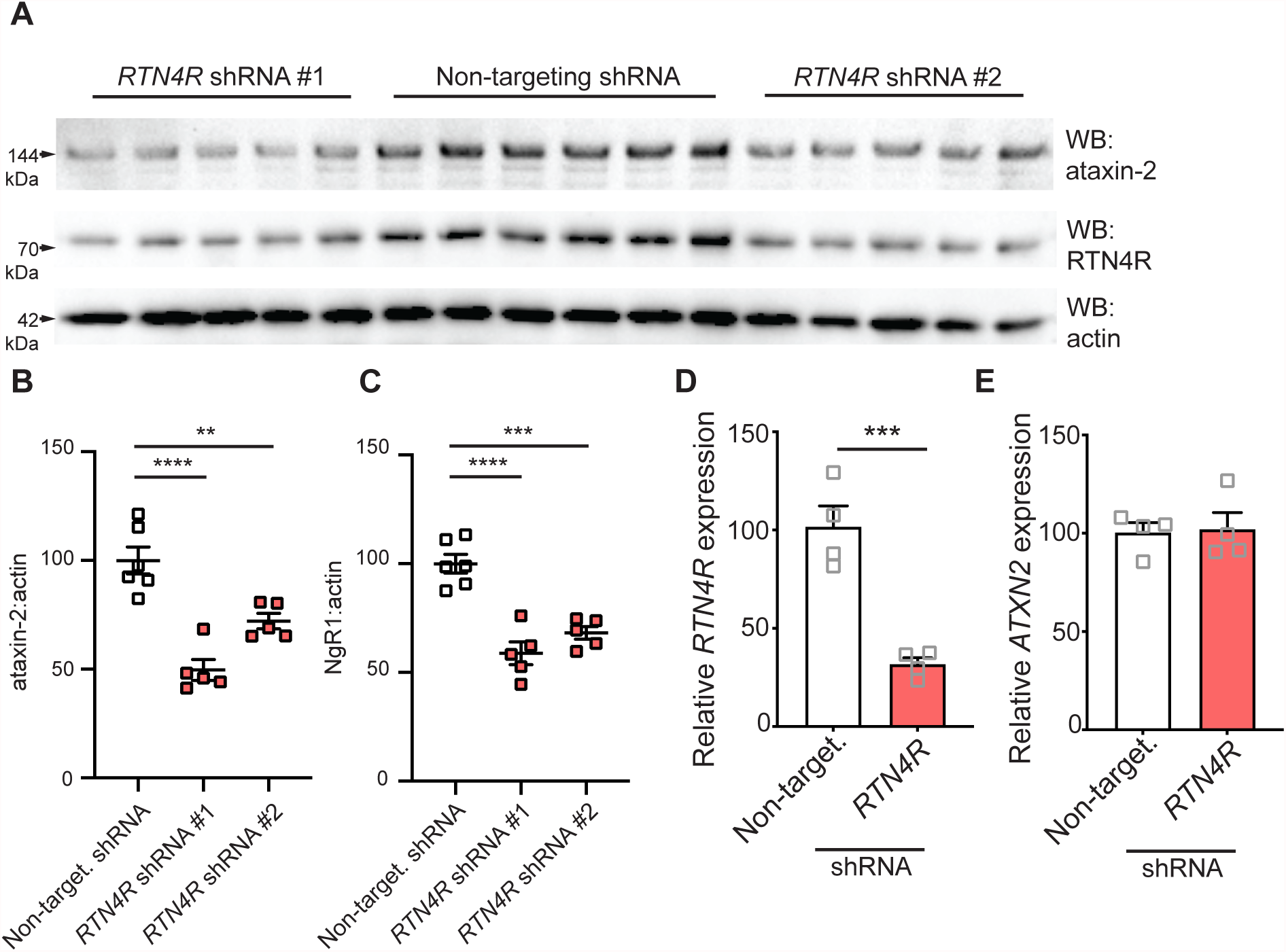
Full immunoblot on neuron lysates treated with two different shRNA constructs targeting *RTN4R* and RT-qPCR on shRNA-treated primary neurons. **(A)** Immunoblot of ataxin-2 and RTN4/NoGo-Receptor levels after 12 days of shRNA treatment in cortical neuron cultures. We used two shRNA constructs targeting different regions of *RTN4R*, along with a non-targeting shRNA construct. Quantification of ataxin-2 **(B)** and RTN4/NoGo-Receptor **(C)** levels. **(D), (E)**, We performed RT-qPCR on RNA from shRNA-treated mouse neurons. We probed for *RTN4R* transcript **(D)** and *ATXN2* transcript **(E)** along with *GAPDH* as a housekeeping gene for normalization. Student’s t-test. **p ≤ 0.01, ***p ≤ 0.001, ****p ≤ 0.0001. Error bars represent ± SEM.

## Acknowledgments

This work was supported by NIH grants R35NS097263(10) (A.D.G.); R35NS097283 (S.M.S.); F32NS116208 (C.M.R.); 5T32NS007280 (G.L.J.); 2T32AG047126-06A1 (T.A.); the Robert Packard Center for ALS Research at Johns Hopkins (A.D.G.); Target ALS (A.D.G.); and the Brain Rejuvenation Project of the Wu Tsai Neurosciences Institute (A.D.G.). G.K. is supported by a fellowship from the Stanford Knight-Hennessy Scholars Program. T.A. is supported by a fellowship from the Takeda Science Foundation. The whole genome siRNA screen was performed with the expertise and resources in the High-Throughput Bioscience Center at Stanford University. Some of the figures were created with BioRender.com.

## Data and Code Availability

The datasets generated during the current study are available from the corresponding author upon request.

